# Major histocompatibility complex class IIB disassortative mate choice in a genetically monogamous seabird

**DOI:** 10.1101/2024.12.19.629314

**Authors:** Lucie Thel, Maxime Pineaux, Léa Ribeiro, Etienne Danchin, Shannon Whelan, Scott A. Hatch, Pierrick Blanchard, Sarah Leclaire

## Abstract

Among species reproducing sexually, mating strategies represent a major component of individual fitness. The major histocompatibility complex (MHC) is an extremely diverse set of genes responsible for immunological recognition and defence against pathogens. Although dissimilarity between mates at the major histocompatibility complex has been proposed to drive mate choice through increased offspring pathogen resistance, evidence is mixed. In addition, explorations of the role of the major histocompatibility complex in other mating strategies, such as divorce, are rare. We investigated whether dissimilarity at the major histocompatibility complex class IIB is associated with mate choice and divorce probability in the genetically monogamous black-legged kittiwake (*Rissa tridactyla*). We found that first-time male breeders, as well as divorced males, were paired with females more dissimilar at the major histocompatibility complex class IIB than expected under random mating. We did not find evidence for mate choice based on major histocompatibility complex class IIB dissimilarity when considering females. In addition, in the studied population, divorce probability was very low compared with other populations and did not significantly vary with the dissimilarity of the pair at the major histocompatibility complex class IIB. Our results pave the way to a better understanding of the complex role of major histocompatibility complex dissimilarity in mating decisions of species displaying mutual choice and biparental care.

## Introduction

Mating strategies are the set of behavioural tactics used by individuals to attract, select, change or retain mates to promote individual fitness. Understanding their driving factors has long been a major topic in evolutionary biology [1,2]. For instance, mate choice has repeatedly been shown to rely on traits that honestly reflect the quantity or quality of resources, the care or the genes that the mate can provide [3–5]. In addition, in several species, individuals preferentially choose mates that are more genetically or behaviourally compatible to avoid inbreeding or to enhance behavioural coordination or task complementation between mates [6–10]. However, individuals may face energetic, time or spatial constraints in choosing mates [11]. Under such circumstances, individuals are forced to choose from a limited pool of potential mates and thus may accept suboptimal partners [12]. A possible response to this limitation in iteroparous species is divorce, defined as one breeding individual changing partner for a new mate while the original mate is still alive [13]. Theoretical studies have suggested that individuals should divorce when pairing with a higher-quality or more compatible mate improves their reproductive success [14]. Accordingly, divorce is common after a reproductive failure [15–17] and usually leads to increased reproductive success [18,19]. Like mate choice, divorce may be associated with the quality of the ornaments of the mate or the behavioural or genetic compatibility of the pair (e.g. [20,21]).

Concerning the genetic component of the pool of heritable factors that can affect mating strategies, the major histocompatibility complex (MHC, called HLA in humans) has been the focus of much research [22–24]. The MHC, one of the most polymorphic sets of genes in vertebrates, encodes cell-surface proteins that are essential for immunological recognition and defence against pathogens. Because each MHC molecule can bind a limited set of antigens [25], higher MHC diversity increases the number of antigens recognized, thereby providing resistance to a wider range of parasites [26,27]. Accordingly, several studies have found that higher MHC diversity increases health, reproductive success and survival [28,29]. Because the MHC allelic composition of the offspring results from the combination of those of the parents, multiple studies have found evidence for mate choice maximizing MHC dissimilarity between mates (e.g. [30–33]). However, high MHC diversity can also increase the risk of autoimmune diseases and reduce the T-cell receptor diversity [34,35]. In a few species, intermediate MHC diversity has thus been shown to be optimum for fitness and to be an important factor in mate choice (e.g. [36–39]). Although there is evidence that the MHC composition or diversity of the parents also influences other reproductive strategies (e.g. cryptic female choice [40]; extra-pair copulation [41]), to our knowledge, a single study on Seychelle warblers (*Acrocephalus sechellensis*) explored the association between parental MHC similarity and divorce rate and found no evidence [42].

Here, we tested whether mate choice and divorce depend on dissimilarity at MHC class-IIB (hereafter MHC-IIB dissimilarity) in the black-legged kittiwake (*Rissa tridactyla*), a long-lived, genetically monogamous seabird with no extra-pair copulation [43]. In black-legged kittiwakes, divorce is fairly common (i.e. 15–50% of the pairs divorce each year) and more frequent among first-time breeders and after reproductive failure [10,44–46]. Coulson [44] suggested, therefore, that some kinds of ‘incompatibility’ between mates result in unsuccessful breeding, thereby providing costs that select for divorce. In black-legged kittiwakes, individuals can recognize MHC-IIB dissimilarity using body odours [47], and MHC-IIB similarity between mates results in reduced offspring MHC-IIB diversity [48], which incurs costs for female chicks such as slower growth, poor tick clearance efficiency and lower survival [49]. We therefore hypothesized that, in this species, mating decisions based on MHC-IIB dissimilarity between mates may have evolved to maximize the MHC-IIB diversity of the offspring. However, the association between MHC diversity and fitness has never been explored in adults in this species, so one cannot exclude that intermediate MHC diversity is optimum when considering lifetime fitness. For all our analyses, we thus also explored the intermediate optimum hypothesis that predicts that individuals should mate with partners with intermediate MHC dissimilarity [36].

First, we tested whether individuals chose their mates according to MHC dissimilarity (Test 1). In several seabirds, including the black-legged kittiwake, young or inexperienced individuals arrive later at the breeding ground than older or more experienced breeders [50–53], which may increase the temporal constraints and limit the pool of potential mates. Inexperienced breeders may also be less efficient in sampling mates and less able to make mating discrimination or to anticipate alternative choices [54]. For instance, in grey mouse lemurs (*Microcebus murinus*), young females in their first mating season do not express mate choice, while experienced females prefer MHC-IIB dissimilar males [55]. The mate choice of first-time breeders may thus be more constrained than that of experienced breeders. We therefore tested the existence of MHC-IIB-dependent mate choice in first-time breeders (Test 1.1) and experienced breeders after a divorce (Test 1.2). Second, if divorcing allows individuals paired with suboptimal mates to optimize the MHC-IIB diversity of their future offspring, we expected the rate of divorce to depend on the MHC-IIB dissimilarity between mates (Test 2). Third, we expected that, after a divorce, the new pair should be more MHC-IIB compatible (maximized or intermediate dissimilarity) than the original pair (Test 3). In black-legged kittiwakes, although both mates participate fairly equally in incubation and chick rearing, sexes slightly differ in energetic investment in reproduction [56,57]. We thus tested these predictions separately in males and females to explore potential sex differences in mating decisions.

## Material and methods

### Study area and population

This study was conducted between 1996 and 2023 in a colony of black-legged kittiwakes nesting in an abandoned U.S. Air Force radar tower on Middleton Island (59°26′ N, 146°20′ W), Gulf of Alaska, located approximately 100 km south of the mainland. Individuals were tagged with a unique combination of coloured and metal bands for long-term monitoring [58]. About 400 nest sites were monitored on the tower each year. All the nests were checked daily to record the identity of the present individual(s). This study was approved by a United States Geological Survey Institutional Animal Care and Use Committee, with permits from the United States Fish and Wildlife Service and the Alaska Department of Fish and Game. For each individual of the study, sex was determined, in order of priority, by: (i) molecular assessment (n = 840 individuals [59]), (ii) behavioural observations (n = 438 individuals [60]), (iii) comparison of the relative size of both members of a pair, the largest bird being considered male (n = 6 individuals), and (iv) when only one individual of a pair was sexed, we attributed the opposite sex to its mate (n = 214 individuals).

Like many populations monitored in the long term, our population has been subjected to several treatments during the study period (e.g. supplemental feeding, brood manipulation, feather clipping, use of anti-copulatory devices). These treatments can affect reproductive success [58,61] and therefore pair stability. To limit any bias (e.g. artificial increase of reproductive success and thus decreased probability of divorce due to supplemental feeding), we excluded pairs that were subjected to experiments from the analysis on the probability of divorce (Test 2).

We defined a pair as two individuals of opposite sex involved in a reproductive event, i.e. attending the same nest and laying at least one egg during a given reproductive season. Individuals usually remain and breed in the same colony during their whole life [44]. We thus defined the first mate choice as the pair formed during the first observed breeding attempt. Each individual was involved in, on average, 4.75 ± 3.74 (s.e.) reproductive events (range = [1–21]; electronic supplementary material, S1) over the course of the study. To study divorce, we focused on individuals that were observed with an identified mate during at least 1 year and that re-mated with a new mate the next year, the original mate being re-observed at least once during the following years [10].

We performed all analyses using the R software [62]. All datasets and codes used in the analyses are available from the Zenodo Repository [63].

### Molecular analyses

We focused on the MHC class II that exhibits stronger selection than the MHC class I in non-passerine birds such as the black-legged kittiwake [64]. Dissimilarity at MHC-IIB was assessed following the methods described in Pineaux et al. [47,49]. Briefly, DNA was extracted from blood with the blood and tissue DNA extraction kit (Qiagen), and a 258 bp fragment of exon 2 of MHC-IIB was amplified using specific primers (forward: 5′-GCACGAGCAGGGTATTTCCA and reverse: 5′-GTTCTGCC ACACACTCACC). Amplicons were then sequenced with an Illumina MiSeq platform, using the 2 × 300 bp protocol (Fasteris SA, Switzerland). Amplicon sequences were analysed with ampliSaS, a three-step pipeline that consists of read demultiplexing, unique sequence clustering and erroneous sequence filtering (substitution errors: 1%; indel errors: 0.001%; minimum frequency with respect to dominant: 33%; minimum amplicon sequence frequency: 2.5% [65]). We calculated the MHC-IIB functional distance between individuals based on amino acid variations in the peptide-binding region (PBR) inferred from Leclaire et al. [66].

We described the chemical binding properties of each amino acid in the PBR with Sandberg’s five physico-chemical descriptors (z-descriptors [67]) and computed Sandberg’s distance matrix between sequences using the DistCalc function of the MHCTools package [68]. Then, we used an unweighted pair group method with arithmetic mean (UPGMA) dendrogram of sequences (hclust function) to represent clusters of functionally similar MHC-IIB sequences (hereafter, ‘supertypes’). The MHC-IIB functional distance between a male and a female was estimated using the tree distance (treedist function of the vegan package [69]), which is found by combining sequences in a common dendrogram and calculating how much of the sum of the branch length is shared and how much is unique. In addition, we calculated the difference in MHC-IIB supertype sharing between mates derived from Strandh et al. [70]:

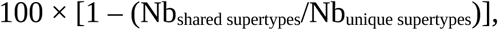

where Nb_shared supertypes_ corresponds to the number of supertypes carried by both the female and the male and Nb_unique supertypes_ corresponds to the total number of unique supertypes possessed by the pair (where common supertypes count as 1). This index thus varies between 0 for individuals sharing all their supertypes and 100 for individuals sharing no supertype. The MHC-IIB functional distance was highly correlated with the difference in MHC-IIB supertype sharing (Spearman correlation test: ρ = 0.79, p < 0.0001, n = 959 pairs; electronic supplementary material, S2).

Mate choice for MHC-IIB dissimilarity may not only be governed by evolutionary processes that specifically target these genes but may also be affected by mechanisms operating at the level of the entire genome (e.g. inbreeding avoidance [71]). To test whether the associations between MHC-IIB dissimilarity and mate choice may be driven by genome-wide relatedness, we used restriction site-associated sequencing (RAD-seq, n = 255 individuals) to estimate genome-wide relatedness between individuals (see detailed methods in electronic supplementary material, S3).

### Statistical analyses

To test the association between mate choice and MHC-IIB dissimilarity (Test 1), we used bootstrapping to generate random pairs, where we let each individual choose randomly 10 000 times between all available opposite-sex mates of the same year, without replacement (i.e. one mate can be selected only once during a given year [72,73]). To test for the maximization hypothesis, we calculated p-values as the proportion of iterations in which the random MHC-IIB dissimilarity mean was greater than the observed MHC-IIB dissimilarity mean, plus the proportion of iterations in which the random MHC-IIB dissimilarity mean was smaller than the symmetrical (relative to the mean of random MHC-IIB dissimilarity means) of the observed MHC-IIB dissimilarity mean (two-tailed test, following the method described in Huchard et al. [55]). To test for the intermediate optimum hypothesis, we used a similar procedure, but instead of comparing the observed and random means of MHC-IIB dissimilarity, we compared the observed and random variance in MHC-IIB dissimilarity (following the method described in Zhang et al. [73]). In the case of a mate choice for intermediate MHC-IIB dissimilarity, we expected the observed and random means of MHC-IIB dissimilarity between partners to be similar. However, because individuals are expected to prefer a mate that allows them to reach the intermediate MHC-IIB dissimilarity optimum, we expected the observed variance in MHC-IIB dissimilarity to be smaller than the random variance. We tested MHC-IIB-based mate choice for first-time breeders (n = 534 females and n = 554 males; Test 1.1) and for experienced breeders after a divorce (n = 113 divorces in females and n = 83 divorces in males; Test 1.2) separately.

To test whether divorce probability varied with the MHC-IIB dissimilarity of the pair during the year n (Test 2), we defined a binary response variable as 1 = divorce between the year n and n + 1, and 0 = no divorce between the year n and n + 1. We tested females (n = 1244) and males (n = 1232) separately. We used generalized linear mixed models (GLMM) with a binomial distribution of error and a logit link (glmer function of the lme4 package [74]). To test for the intermediate optimum hypothesis, we included the quadratic term of the MHC-IIB dissimilarity [75]. If this variable did not have a significant effect (i.e. p > 0.05), we removed the quadratic term and then tested for the maximization hypothesis with the linear term of the MHC-IIB dissimilarity. The reproductive outcome is also known to influence the probability of divorce in black-legged kittiwakes [10,46]. We thus defined the reproductive outcome as a binary variable recording the presence (i.e. ‘success’) or absence (i.e. ‘failure’) of at least one chick fledged at the end of the reproductive season and included this variable as an additive effect. In the models, we set the nAGQ argument (number of points per axis for evaluating the adaptive Gauss–Hermite approximation to the log-likelihood) of the glmer function to 0 to help for convergence, when necessary. We tested the effect of individual identity and year as random intercepts with the ranef function of the lme4 package [74] and removed these variables when they had no effect. We removed pairs that were subjected to experiments from this analysis. As we used GLMM with binomial data, we provided the log odds (and its standard error), which is equivalent to the coefficient of the slope in a linear model (i.e. estimate in the model summary). It corresponds to the probability of an event on the log scale, which is the transformed scale to satisfy the assumptions of a linear model for GLMM with binomial data. For convenience, we also provided the odds ratio, which corresponds to the exponential of the log odds. An odds ratio of 1 means no effect, whereas an odds ratio >1 (<1, respectively) corresponds to an increase (decrease, respectively) of the response variable when the explanatory variable (here, reproductive outcome or MHC-IIB dissimilarity) increases by 1 unit. We calculated the confidence intervals from the model on the fixed effects using the predictInterval function of the merTools package (n iterations = 10 000, [76]).

To test whether, after divorce, the new mates were more MHC-IIB dissimilar than the original mates (i.e. the maximization hypothesis), we used linear mixed models (LMM; Test 3). The response variable was the MHC-IIB dissimilarity gain, i.e. the difference between the MHC-IIB dissimilarity to the original mates and the MHC-IIB dissimilarity to the new mates. Using the gain in a linear modelling framework allowed us to include individual and year as random effects. The MHC-IIB composition of the individuals available in the population might vary with year, thereby potentially affecting the ability to find MHC-IIB dissimilar mates. The use of random effects allowed us to account for this potential source of variability. We tested the effect of individual identity and year as random intercepts with the ranef function of the lme4 package [74] and removed these variables when they had no effect. We tested the intermediate optimum hypothesis by comparing the variance of the MHC-IIB dissimilarity between the pool of original pairs and the pool of new pairs with a Fisher exact test (or its non-parametric equivalent, the Levene test, when necessary). In the case of mate choice following the intermediate hypothesis, we expected the variance of the MHC-IIB dissimilarity in the new pairs to be lower than the variance of the MHC-IIB dissimilarity in the original pairs.

### Power analyses

We conducted test-specific power analyses to assess the robustness of our results. When studying the MHC-IIB dissimilarity with the new mates compared with random mates (Tests 1.1 and 1.2), we evaluated the likelihood of error as detailed in Hoover & Nevitt [72] and Hoover et al. [77]. For example, for males involved in divorces, we resampled between 10 and 83 males (the actual number of individuals in this dataset) and conducted the same analysis as described previously (nboots = 500 instead of 10 000 for reasons of computational limitations). We then assigned a ‘1’ to a significant result and a ‘0’ to a non-significant result for every sample size tested and used a GLM with a binomial distribution of error and a logit link (glmer function of the lme4 package [74]) to describe the relationship between sample size and likelihood of error. We also provided the effect size measured as the difference between the observed MHC-IIB dissimilarity mean (respectively variance) and the random MHC-IIB dissimilarity mean (respectively variance).

When assessing the probability of divorce according to MHC-IIB dissimilarity (Test 2), our dataset was large enough (>1000 individuals) but largely unbalanced between faithful and divorcing pairs (approximately 20 times less divorced than faithful pairs in both females and males). We thus randomly resampled the faithful pairs to obtain the same sample size as for divorced pairs and adjusted the same models (nboots = 1000) as described previously on the balanced dataset using GLMs. We then estimated the probability of obtaining a significant effect for each explanatory variable (i.e. quadratic and linear terms of MHC-IIB dissimilarity, and reproductive outcome).

When assessing the gain in MHC-IIB dissimilarity between the original and the new pairs after a divorce (Test 3), we calculated the effect sizes for our analysis using Cohen’s d statistic [78].

## Results

### Mate choice in first-time breeders (Test 1.1)

Most of the first-time breeders mated with first-time breeders (62% when considering males and 65% when considering females). For their first breeding attempt, males tended to mate with females that had a significantly greater difference in supertype sharing than expected under random mating (p = 0.05, effect size = 1.61, figure 1b; electronic supplementary material, table S4.1). The probability of finding a significant result with the available sample size (n = 554 mate choice events) was 0.40 (electronic supplementary material, table S4.1 and figure S4.1b). This effect was, however, not significant when considering the functional distance between mates (p = 0.30, effect size = 0.07; probability of finding a significant result = 0.03; electronic supplementary material, figure S4.2c and S4.2d). Females and their first mates did not show significantly greater difference in supertype sharing or greater functional distance than expected under random mating (functional distance: p = 0.89, effect size = −0.01, electronic supplementary material, figure S4.2a; difference in supertype sharing: p = 0.72, effect size = 0.31, figure 1a; electronic supplementary material, table S4.1). Neither females (functional distance: p = 0.98; difference in supertype sharing: p = 0.88) nor males (functional distance: p = 0.75; difference in supertype sharing: p = 0.49) mated with significantly more intermediately dissimilar mates than expected under random mating (electronic supplementary material, table S4.1 and figures S4.3 and S4.4).

**Figure 1.**
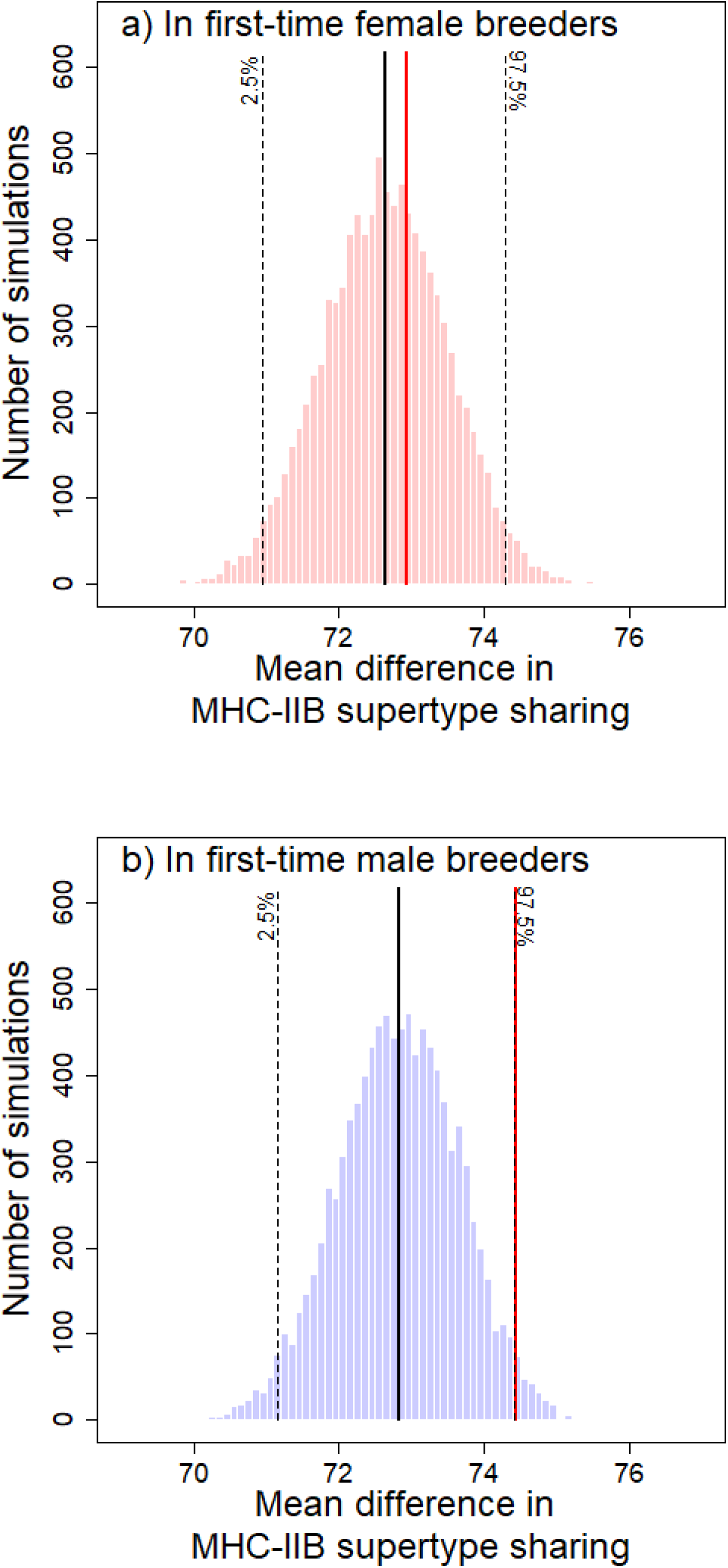
Distribution of 10 000 means of MHC-IIB dissimilarity (as measured by the difference in MHC-IIB supertype sharing) between first-time breeders and their mates, calculated from bootstrap randomizations of potential pairs (see §2 for details), when considering (a) females and (b) males. The red vertical line is the observed mean of MHC-IIB dissimilarity. The black vertical line is the mean of the simulated distribution, and the black dashed lines are the 2.5 and 97.5% quantiles of the simulated distribution showing the 95% confidence intervals. See electronic supplementary material, figure S4.1, for a representation of the likelihood of a type II error according to the sample size.

### Probability of divorce (Test 2)

In both sexes, the probability of divorce was significantly lower after a reproductive success than after a reproductive failure (all p ≤ 0.01; probability of finding a significant result with a balanced dataset >0.80; electronic supplementary material, table S5.1 and figure S5.1). However, the probability of divorce did not significantly vary with the MHC-IIB dissimilarity of the pair (electronic supplementary material, figure S5.2), when considering either females (linear term: functional distance: p = 0.58 and difference in supertype sharing: p = 0.74; quadratic term: functional distance: p = 0.18 and difference in supertype sharing: p = 0.46; electronic supplementary material, table S5.1) or males (linear term: functional distance: p = 0.68 and difference in supertype sharing: p = 0.54; quadratic term: functional distance: p = 0.64 and difference in supertype sharing: p = 0.81; electronic supplementary material, table S5.1). The probability of finding a significant effect of MHC-IIB dissimilarity on the probability of divorce was consistently <0.20 with a balanced dataset (electronic supplementary material, table S5.1).

### Mate choice in divorced individuals (Tests 3 and 1.2)

Divorced individuals re-mated mainly with experienced breeders (63% when considering males and 71% when considering females). The difference in MHC-IIB dissimilarity between the new pair and the original pair was not different from 0, either in females or males (Test 3 – maximization hypothesis: all p > 0.10; electronic supplementary material, table S6.1). The effect sizes revealed by Cohen’s d statistics were very low (all Cohen’s d < 0.20; electronic supplementary material, table S6.1 and figure S6.1).

However, after a divorce, males were paired with females, which had a significantly greater difference in supertype sharing to them than expected under random mating (Test 1.2: p = 0.04, effect size = 4.52; probability of finding a significant result = 0.78; figure 2b and electronic supplementary material, figure S7.1b and table S7.1). Females were not re-mated with males, which had a significantly greater difference in supertype sharing than expected under random mating (Test 1.2: p = 0.06, effect size = 3.58; probability of finding a significant result = 0.23; figure 2a and electronic supplementary material, figure S7.1a and table S7.1). These effects were not significant when considering functional distance (males: p = 0.09, effect size = 0.31 and females: p = 0.62, effect size = 0.08; electronic supplementary material, table S7.1 and figure S7.2).

**Figure 2.**
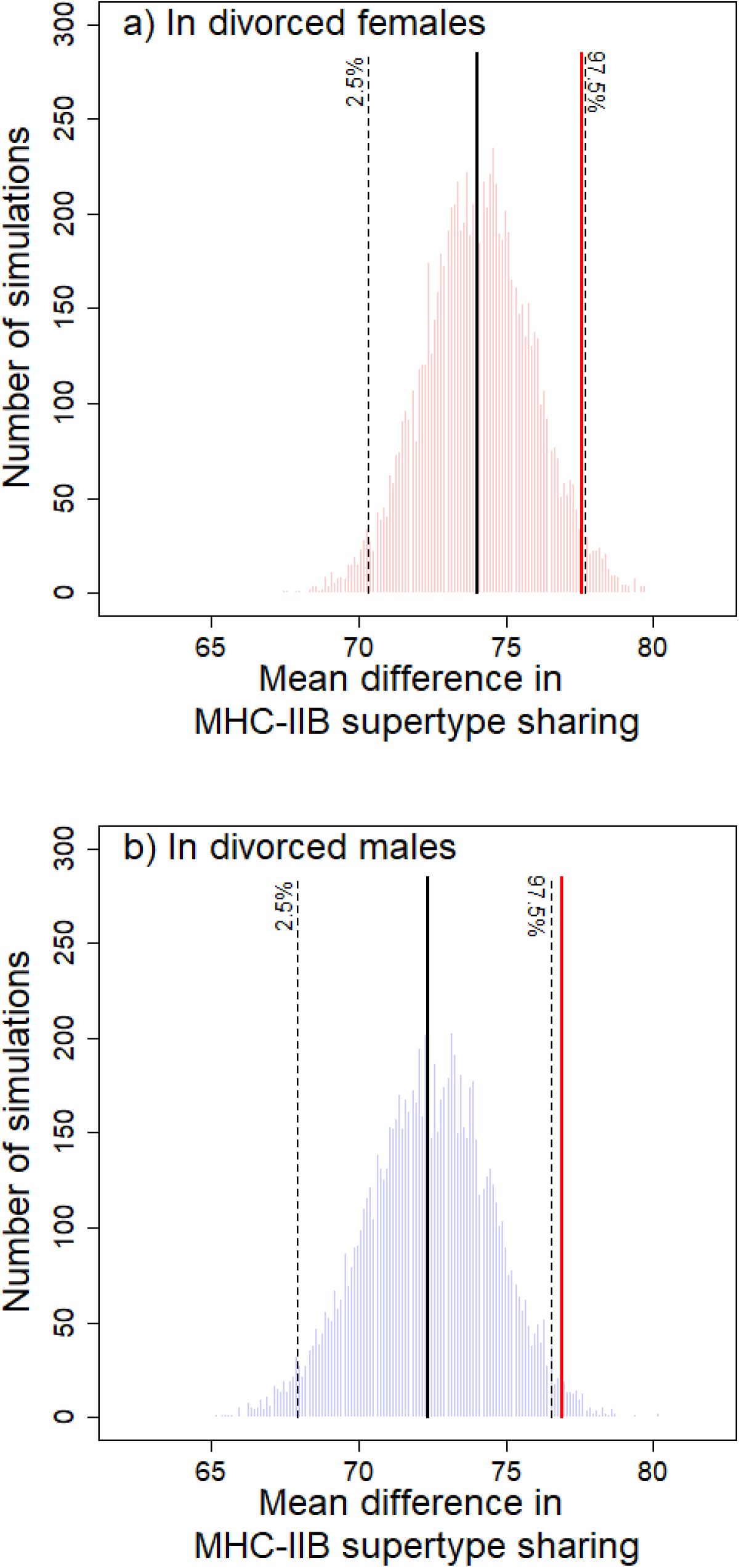
Distribution of 10 000 means of MHC-IIB dissimilarity (as measured by the difference in MHC-IIB supertype sharing) between divorced breeders and their new mates, calculated from bootstrap randomizations of potential pairs (see §2 for details), when considering (a) females and (b) males. The red vertical line is the observed mean of the MHC-IIB dissimilarity. The black vertical line is the mean of the simulated distribution, and the black dashed lines are the 2.5 and 97.5% quantiles of the simulated distribution showing the 95% confidence intervals. See electronic supplementary material, figure S7.1, for a representation of the likelihood of a type II error according to the sample size.

When considering either males or females, after a divorce, the new pairs were not more intermediately dissimilar than the original pairs (Test 3 – intermediate hypothesis: all p > 0.10; electronic supplementary material, table S6.1) or than random pairs (Test 1.2 – intermediate hypothesis: all p > 0.10; electronic supplementary material, table S7.1).

### MHC-IIB dissimilarity and genome-wide relatedness

We found that genome-wide relatedness was correlated neither with functional MHC-IIB similarity (Mantel test: r = 0.04, p = 0.09; electronic supplementary material, figure S3.1) nor with difference in MHC-IIB supertype sharing (Mantel test: r = 0.03, p = 0.10; electronic supplementary material, figure S3.2).

## Discussion

In this study, we tested whether mating decisions (i.e. mate choice and divorce) of black-legged kittiwakes vary with the MHC-IIB dissimilarity between mates. We found that first-time male breeders (Test 1.1) as well as divorced males (and to a lower extent, divorced females; Test 1.2) were paired with mates more MHC-IIB dissimilar to them than expected under random mating. However, we did not detect an association between the MHC-IIB dissimilarity of the pair and the probability of divorce.

First-time male breeders, as well as divorced males, were more MHC-IIB dissimilar to their female partners than expected under random mating. These results may suggest active mate choice for MHC-IIB dissimilarity. Like several other species (e.g. [26,79–81]), including birds [82,83], black-legged kittiwakes are able to recognize MHC-IIB dissimilarity using body odours [47]. In this species, the recognition of MHC-IIB dissimilarity might have evolved as a mechanism to select MHC-IIB dissimilar mates to produce offspring with high MHC-IIB diversity [48], and thus with higher early-life performance [49]. Similarly, in the domestic cat (*Felis catus*), mating with MHC-IIB dissimilar mates increases offspring survival and thus probably parental fitness [84].

In contrast to results when considering males, we did not detect MHC-IIB disassortative mate choice when considering first-time female breeders. In addition, we detected only a tendency for MHC-IIB disassortative mate choice when considering divorced females (p = 0.06, with 23% likelihood of obtaining a significant result). Our analysis design, however, does not clearly allow us to determine whether the MHC-IIB disassortative mate choice detected when considering males is driven by males’ choice or females’ choice. For instance, an MHC-IIB disassortative mate choice detected in divorced males may suggest that divorced males preferentially choose MHC-IIB dissimilar females or that females preferentially select MHC-IIB dissimilar males when they have to choose among divorced males. Similarly, a lack of MHC-IIB disassortative mate choice detected when considering first-time female breeders may be due to first-time female breeders not choosing male partners based on MHC-IIB dissimilarity or to males not choosing MHC-IIB dissimilar female partners when they have to choose among first-time female breeders.

The contrasting results between males and females might, however, illustrate sex-specific mating strategies. For instance, in Leach’s storm-petrels (*Oceanodroma leucorhoa*), while males show MHC-IIB disassortative mate choice, females’ choice does not differ from random mating [77]. In long-lived monomorphic species in which males and females equally share parental care, such as the black-legged kittiwake [85–87], mate choice has been suggested to be mutual [12]. However, the trade-off between the benefits obtained through MHC disassortative mate choice and those obtained through mate choice based on the intrinsic quality of the partner, such as its ability to provide good genes [88,89], good parental care [90] or good-quality territory [91], can vary between individuals [88,92], including between sexes. It may thus result in males and females having different preference criteria for mate choice [93,94]. Sex differences in the relative influence of MHC dissimilarity versus other traits on mating preference might be due to sex-specific variability in either of these traits or to sex-specific variations in the direct benefits of these other traits to the chooser. For instance, in mice, MHC dissimilarity is a more important criterion for mate choice than scent-marking rate, but only when variability in MHC dissimilarity between mates is large [92]. In a cichlid fish (*Pelvicachromis taeniatus*), body size is a more important criterion for mate choice than genetic compatibility for males than for females, which may be due to the crucial role of female body size on fecundity [8].

When using Monte Carlo simulations to test mate choice, Hoover & Nevitt [72] showed that a sample size of 500 pairs may be necessary to confidently reject the null hypothesis when the effect size is small (see also [75]), small effect sizes being common when studying MHC-based mate choice [75]. In the present study, the sample size of first-time breeding males was ca 500 individuals, and we found a p = 0.05 in males with a 60% likelihood of finding a non-significant result. This suggests at best a low effect size for mate choice based on MHC-IIB dissimilarity in first-time breeders. In contrast, the sample size of divorced males was considerably smaller (i.e. only 83 males), but we found a significant MHC-IIB disassortative mate choice (p = 0.04) with a much larger effect size in divorced individuals than in first-time breeders (compare effect sizes in electronic supplementary material, tables S4.1 and S7.1). In several species, the level of mate selectivity is influenced by age [55,95–97]. In black-legged kittiwakes, higher selectivity in divorced individuals compared with first-time breeders may be related to their earlier arrival on the nesting site [50], which may increase both decision time and the pool of potential partners and thereby mating decision accuracy [98]. However, like for the sex-specific variations detected in the mate choice analysis, we cannot determine whether the higher selectivity for MHC-IIB dissimilar partners detected when considering divorcees than when considering first-time breeders is due to divorced individuals being more selective in choosing MHC-IIB dissimilar males or individuals being more selective in choosing MHC-IIB dissimilar mates when they have to choose among divorced partners. For instance, in bluethroats (*Luscinia svecica*), females seem to select males based on MHC-related traits only when paired with young males [39]. In black-legged kittiwakes, in the context of a balance between selecting mates for MHC-IIB dissimilarity versus other individual quality traits, the lower inter-individual variability in individual quality in experienced breeders than in first-time breeders [99] may explain the higher selectivity for MHC-IIB dissimilarity when the pool of potential partners is divorcees. Clearly, while our study provides invaluable insight into the MHC-IIB disassortative mating strategy of black-legged kittiwakes, further studies are needed to determine which sex and age groups are more selective or selected based on MHC-IIB dissimilarity, and why.

In accordance with previous studies in black-legged kittiwakes [10,45,46,100], we found that the probability of divorce was greater in pairs that failed to fledge a chick than in pairs that reproduced successfully. However, it does not significantly vary with the MHC-IIB dissimilarity between mates. Also, after a divorce, individuals do not mate with partners more MHC IIB dissimilar to them than their original partners. Together, these results suggest that in black-legged kittiwakes, MHC-IIB dissimilarity between the male and the female of the pair is not a driving factor of divorce. In the present study, the proportion of divorces was much lower than the proportions observed in other populations of black-legged kittiwakes (annual mean ± s.d. = 4.4 ± 2.6% in our study versus 25% in [44]; between 13 and 47% in [45]; 19.1% on average in [46]). On Middleton Island, numerous black-legged kittiwakes nest on other man-made structures than the tower [58]. These sites were not monitored, except for occasional demographic surveys for certain years. It may, therefore, have limited our ability to observe birds that changed nesting sites between breeding seasons and thus to detect all divorces, leading to a low sample size that might have affected our results and limited our ability to detect a significant effect.

We did not find any support for the intermediate optimum hypothesis in mating decisions in black-legged kittiwakes. This result is consistent with Pineaux et al. [49], where no benefits of intermediate MHC-IIB diversity were detected in black-legged kittiwake chicks. In this species, the benefits of a larger MHC-IIB repertoire associated with increased resistance to a wider range of pathogens may outcompete the costs associated with increased risks of autoimmune diseases and reduced T-cell receptor diversity [34,35].

A previous study on the same population of black-legged kittiwakes showed that pairs have a higher genetic dissimilarity (as measured by microsatellites) than expected under random mating and that the resulting increased heterozygosity of their chicks leads to higher fitness [101]. Besides increasing the resistance to pathogens of the offspring, MHC-IIB disassortative mate choice may limit inbreeding [80,102]. In the present study, we detected MHC-based mate choice when MHC-IIB dissimilarity was measured based on difference in supertype sharing, but not when based on functional distance. Similarly, in blue petrels (*Halobaena caerulea*), yellow-rumped flycatchers (*Ficedula zanthopygia*) and little auks (*Alle alle*), evidence for MHC-based mating strategies is stronger when considering supertype or allele sharing than when considering functional distances [70,103,104] (but see [55] for the opposite pattern). Understanding whether the difference detected between the two indices illustrates that MHC-based mate choice is related to inbreeding avoidance in this species requires a larger number of individuals with genome-wide data.

In conclusion, we found evidence that black-legged kittiwakes choose partners based on MHC-IIB dissimilarity. This finding supports the role of the benefits associated with high MHC-IIB dissimilarity in the evolution of mating decisions in this population. It is consistent with a previous study showing that, in this species, sex-allocation strategy varies with MHC-IIB dissimilarity between parents [48]. In addition, our study suggests that mate choice for MHC-IIB dissimilar partners varies with sex and age or experience of the chooser or the selected mate. When the relative benefits of MHC disassortative mate choice vary with sex and mate choice is mutual, a sexual conflict over mating strategies can arise and may select for a rich repertoire of sex-specific behavioural adaptations [8,105]. Our study paves the road for further studies of sex-specific mate choice in species with biparental care, a common trait in birds [106].

## Declarations

### Ethics

Research activities on Middleton Island, AK, were permitted by the United State Geological Survey (Banding Permit 23910) and the McGill Animal Care Committee (animal use protocol 2016–7814 and precursors).

### Data accessibility

Data available from the Zenodo Repository [63]. Supplementary material is available online [107].

### Declaration of AI use

We have not used AI-assisted technologies in creating this article.

### Authors’ contributions

L.T.: conceptualization, formal analysis, investigation, methodology, visualization, writing—original draft, writing—review and editing; M.P.: conceptualization, data curation, writing—review and editing; L.R.: data curation; E.D.: funding acquisition, project administration, writing—review and editing; S.W.: project administration; S.A.H.: funding acquisition, project administration, supervision; P.B.: conceptualization, data curation, writing— review and editing; S.L.: conceptualization, data curation, formal analysis, funding acquisition, methodology, project administration, resources, supervision, visualization, writing—original draft, writing—review and editing. All authors gave final approval for publication and agreed to be held accountable for the work performed therein.

## Acknowledgements.

We thank the Middleton fieldworkers who collected data, as well as the undergraduate students who helped with molecular analyses.

