## supporting information for "Major histocompatibility complex class IIB disassortative mate choice in a genetically monogamous seabird"

### Authors details

[1] Université de Toulouse, Toulouse INP, CNRS, IRD, CRBE, Toulouse, France.

[2] Department of Conservation Management, Faculty of Science, Nelson Mandela University, George, South Africa.

[3] Centre de Recherches sur la Cognition Animale, Centre de Biologie Intégrative, UMR 5169 (CNRS, Université de Toulouse Midi-Pyrénées), Toulouse, France.

[4] Institute for Seabird Research and Conservation, Anchorage, Alaska, USA.

DOI : https://doi.org/10.1098/rspb, Proceedings of the Royal Society B

### Supporting information

**Supporting information 1**: Number of reproductive events per individual (n = 3064 reproductive events) in the population of black-legged kittiwake (*Rissa tridactyla*) nesting in Middleton Island during the study period (1996-2023).

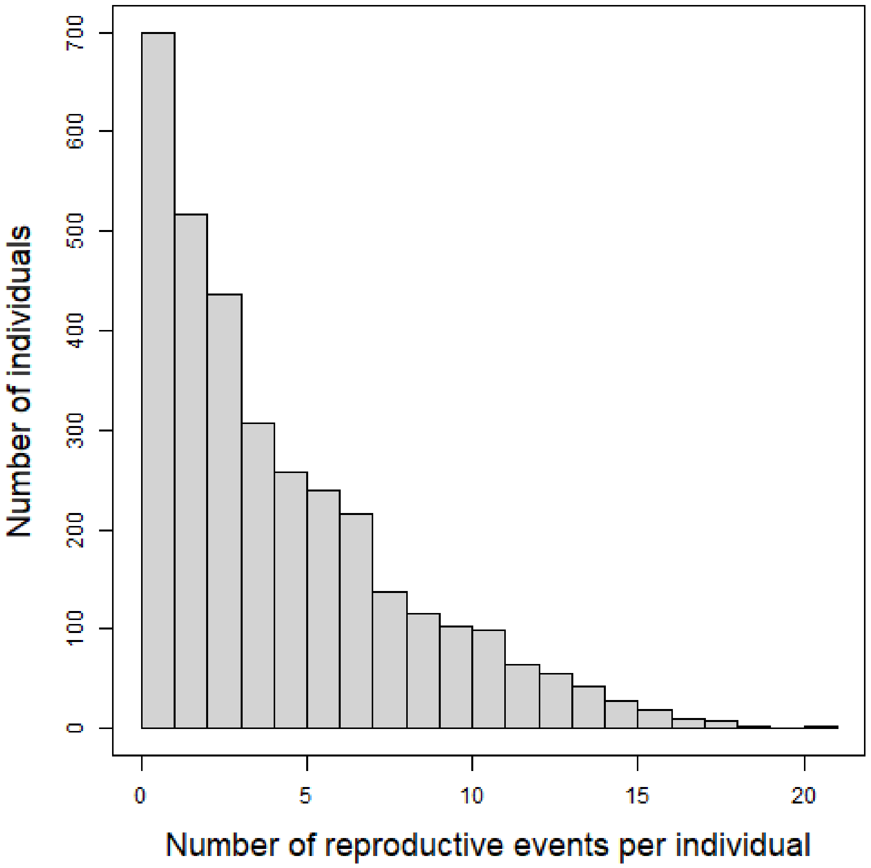

**Supporting information 2**: Associations between MHC-IIB functional distance and difference in MHC-IIB supertype sharing among breeding pairs (n = 959 pairs) of black-legged kittiwake (*Rissa tridactyla*) nesting in Middleton Island during the study period (1996-2023).

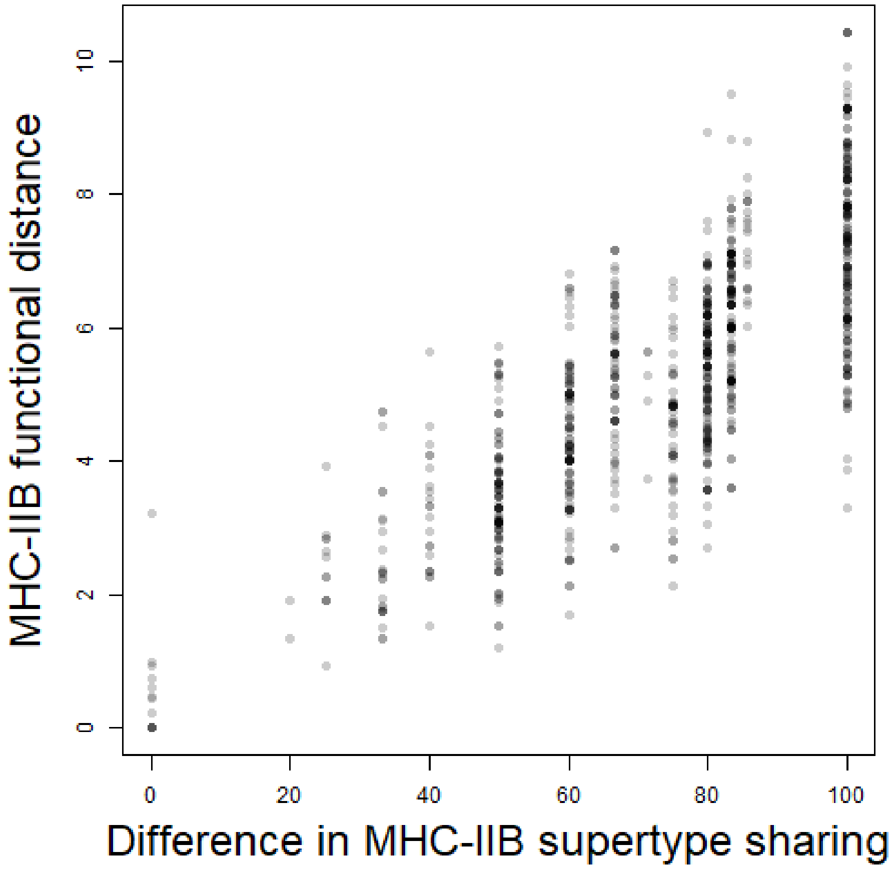

**Supporting information 3**: Associations between MHC-IIB dissimilarity and genome-wide relatedness.

For 255 individuals, DNA was extracted from frozen red blood cells or blood stored in ethanol, with the Blood and Tissue DNA Extraction kit (Qiagen). Six individuals were run in duplicates. DNA was digested using Sbf I restriction enzyme (CCTGCAGG). Samples were sequenced on an Illumina NovaSeq 6000in two sequencing runs. Reads were demultiplexed and cleaned using process_radtags from Stacks, and PCR duplicates were removed using clone_filter from Stacks. The reads were mapped to the reference genome bRisTri1.patW.cur.20221130 (GCF_028500815.1; Sozzoni et al. 2023) with BWA using default settings (Li and Durbin 2009). To generate a high-quality dataset with minimal missing data, we then used PLINK to extract polymorphic SNPs and to retain loci with a genotyping rate of over 90 %, minor allele frequency (MAF) > 0.01 and that conformed to Hardy-Weinberg equilibrium with a p-value threshold of 0.001. One of the two runs had lower average sequencing depth than the other. To ensure comparable locus representation across runs, we therefore only retained loci present in at least 90 % of individuals from the lower-depth run, thereby standardizing data quality and minimizing potential biases introduced by differential sequencing coverage. The number of SNPs was mean ± SE = 11003 ± 33 SNPs, min = 6273 SNPs and max = 11273 SNPs. Relatedness between individuals was calculated using the *relatedness2* function of the *vcftools* package (Danecek et al. 2011), which is based on the methods of Manichaikul et al. (2010). Relatedness between duplicates was mean ± SE = 0.44 ± 0.02, min = 0.37 and max = 0.49, as expected by relatedness > 0.354 being duplicates (Manichaikul et al. 2010). Correlation between genome-wide relatedness and MHC-IIB similarity between individuals (shared supertypes and functional similarity) was tested using Mantel tests and 5000 permutations. We found that genome-wide relatedness was correlated neither with functional MHC-IIB similarity (Mantel test: r = 0.04, P = 0.09; Figure S3.1), nor with MHC-IIB supertype sharing (Mantel test: r = 0.03, P = 0.10; Figure S3.2).

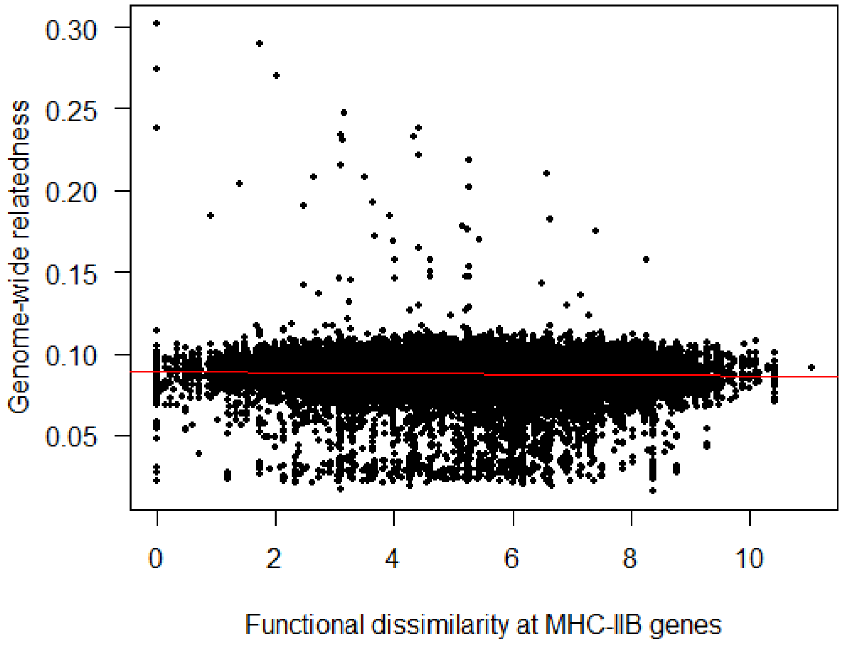


Figure S3.1: Relationship between MHC-IIB functional dissimilarity and genome-wide relatedness. The red line is the regression line between the two variables.

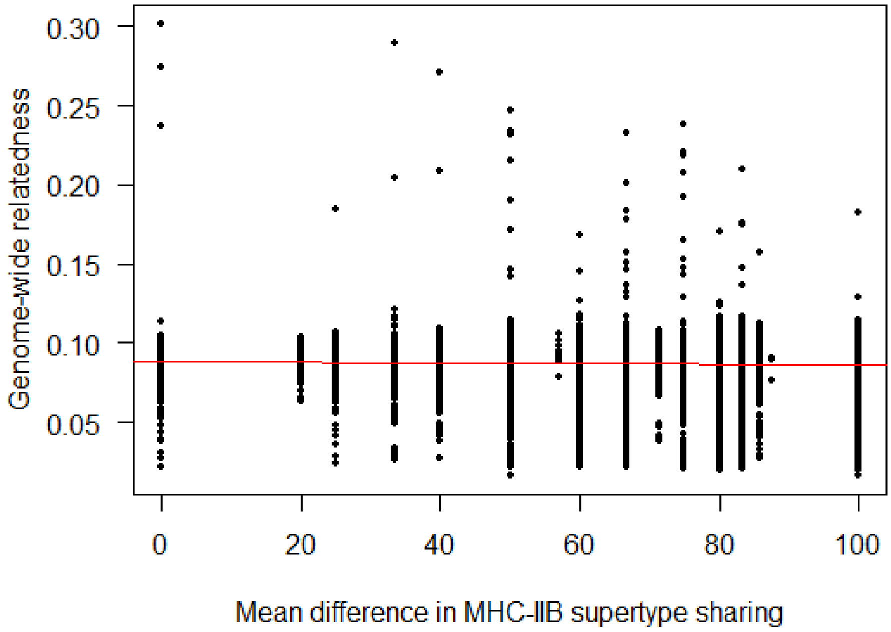


Figure S3.2: Relationship between the difference in MHC-IIB supertype sharing and genome-wide relatedness. The red line is the regression line between the two variables. Pairs with [0.177; 0.354] relatedness are suggested to be first degree relatives, and pairs with [0.0884; 0.177] are suggested to be second-degree relatives (Manichaikul et al. 2010).

**Supporting information 4**: Complementary figures - MHC-IIB based mate choice in first-time breeders (Test 1.1)

Table S4.1: Statistical significance (column "P") of the difference between the observed MHC-IIB dissimilarity (column "Observed value") between the breeding individuals (either females or males, column "Sex of the individual") and thier mates, and the random MHC-IIB dissimilarity mean (column "Random estimate", with 95 % confidence intervals) (Test 1.1). The maximisation hypothesis and the intermediate optimum hypothesis were tested (column "Hypothesis") for both MHC-IIB functional distance and difference in MHC-IIB supertype sharing (column "MHC dissimilarity index"). See main text for details about the "Probability of significant result" and "Effect size" columns.

| **Hypothesis** | **MHC dissimilarity index** | **Sex of the individual** | **P** | **Observed value** | **Random estimate** | **Effect size** | **Probability of significant result** | **Sample size** |
|---|---|---|---|---|---|---|---|---|
| intermediate | functional | female | 0.98 | 3.34 | 3.33 [2.97; 3.71] | 0.01 | 0.00 | 534 |
| maximisation | functional | female | 0.89 | 5.28 | 5.29 [5.15; 5.43] | -0.01 | 0.00 | 534 |
| intermediate | functional | male | 0.75 | 3.40 | 3.34 [3.00; 3.71] | 0.06 | 0.00 | 554 |
| maximisation | functional | male | 0.30 | 5.36 | 5.28 [5.15; 5.42] | 0.07 | 0.03 | 554 |
| intermediate | supertype | female | 0.88 | 452.02 | 447.14 [387.06; 511.37] | 4.88 | 0.00 | 534 |
| maximisation | supertype | female | 0.72 | 72.94 | 72.63 [70.94; 74.29] | 0.31 | 0.00 | 534 |
| intermediate | supertype | male | 0.49 | 419.72 | 440.91 [382.10; 502.91] | -21.19 | 0.00 | 554 |
| maximisation | supertype | male | 0.05* | 74.43 | 72.82 [71.17; 74.41] | 1.61 | 0.40 | 554 |

*: significance to the level alpha = 0.05.

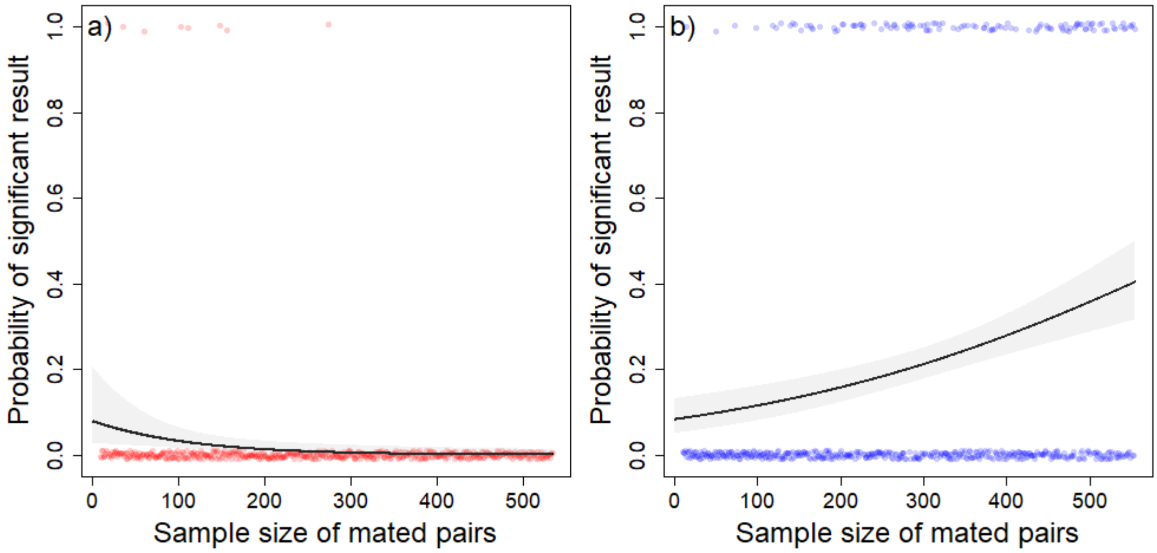


Figure S4.1: Probability of obtaining a significant result according to the sample size when testing the difference in MHC-IIB supertype sharing between first-time breeders and their mates (cf. Figure 1 in the main text), when considering **a)** females, and **b)** males. The dots represent the raw observations from the simulations at each sample size, the line represents the predictions of the model, the shaded area represents the 95 % confidence intervals around the predictions of the model.

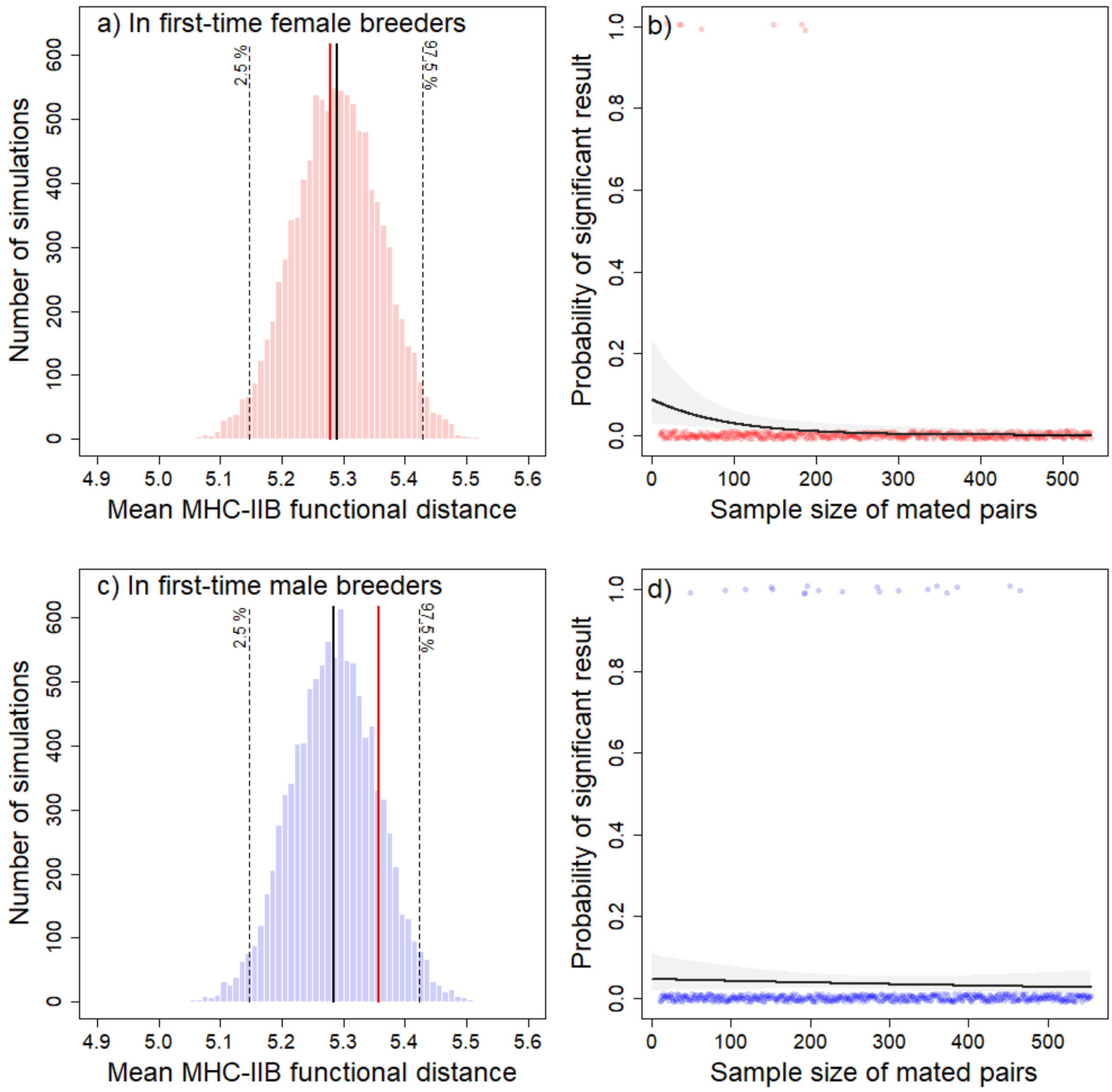


Figure S4.2: **a-c)** Distribution of 10000 means of MHC-IIB dissimilarity (as measured by the MHC-IIB functional distance) between first-time breeders and their mates, calculated from bootstrap randomisations of potential pairs (see Materials and Methods section for details), when considering a) females, and c) males. The red vertical line is the observed mean of MHC-IIB dissimilarity. The black vertical line is the mean of the simulated distribution, and the black dashed lines are the 2.5 and 97.5 % quantiles of the simulated distribution showing the 95 % confidence intervals. **b-d)** Probability of obtaining a significant result according to the sample size, when considering b) females, and d) males. The dots represent the raw observations from the simulations at each sample size, the line represents the predictions of the model, the shaded area represents the 95 % confidence intervals around the predictions of the model.

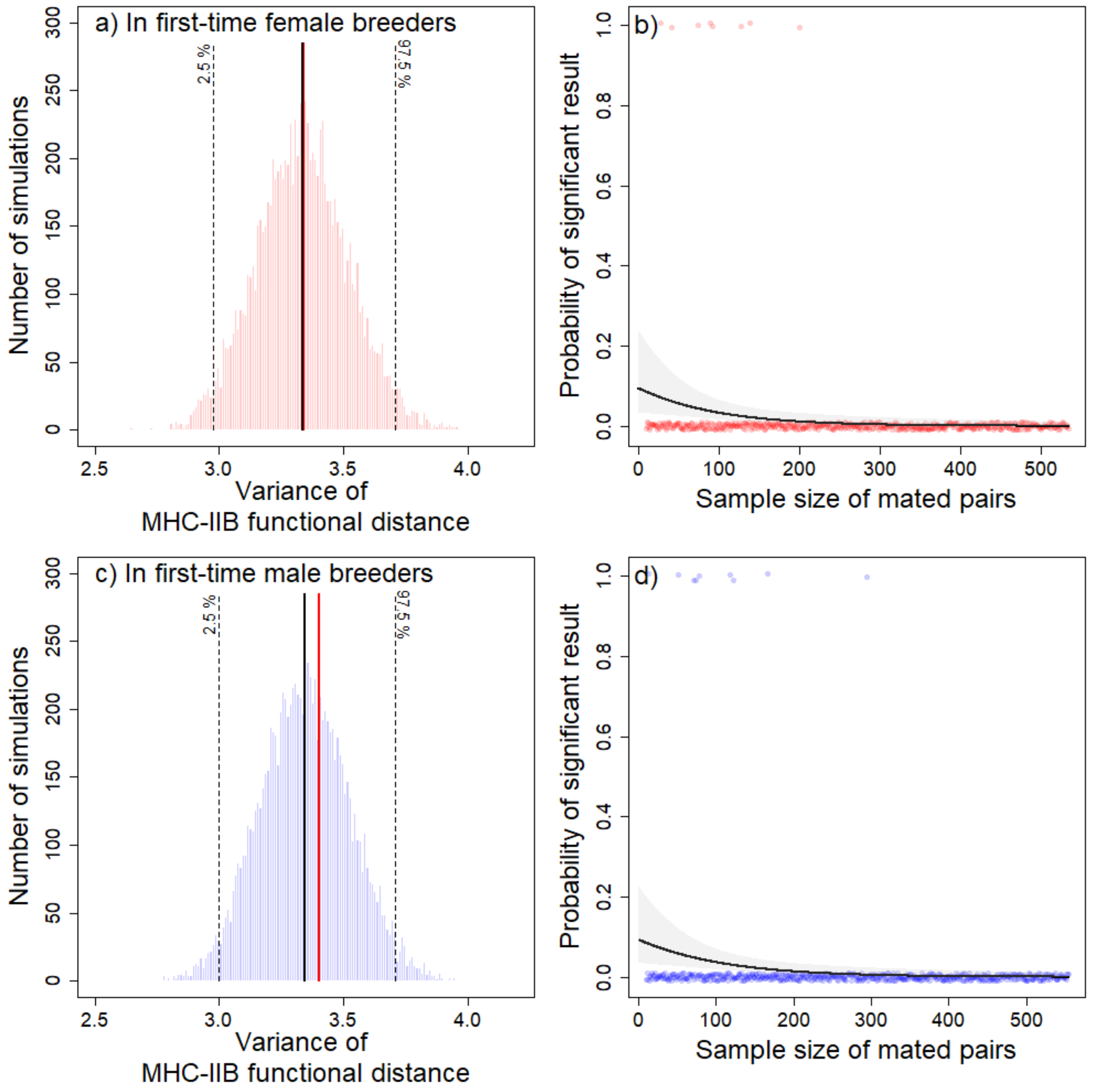


Figure S4.3: **a-c)** Distribution of 10000 variances of MHC-IIB dissimilarity (as measured by the MHC-IIB functional distance) between first-time breeders and their mates, calculated from bootstrap randomisations of potential pairs (see Materials and Methods section for details), when considering a) females, and c) males. The red vertical line is the observed variance of MHC-IIB dissimilarity. The black vertical line is the mean of the simulated distribution, and the black dashed lines are the 2.5 and 97.5 % quantiles of the simulated distribution showing the 95 % confidence intervals. **b-d)** Probability of obtaining a significant result according to the sample size, when considering b) females, and d) males. The dots represent the raw observations from the simulations at each sample size, the line represents the predictions of the model, the shaded area represents the 95 % confidence intervals around the predictions of the model.

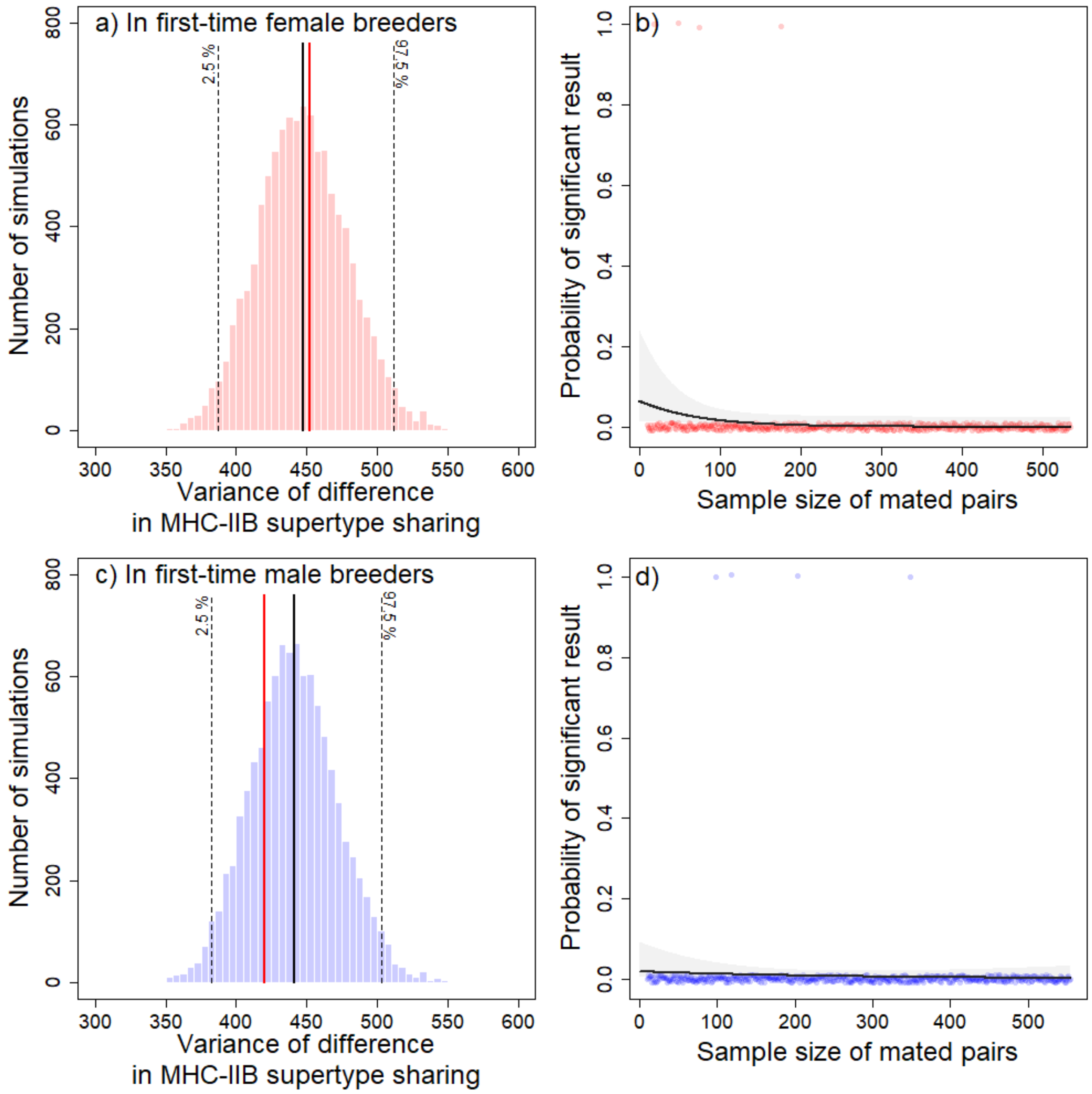


Figure S4.4: **a-c)** Distribution of 10000 variances of MHC-IIB dissimilarity (as measured by the difference in MHC-IIB supertype sharing) between first-time breeders and their mates, calculated from bootstrap randomisations of potential pairs (see Materials and Methods section for details), when considering a) females, and c) males. The red vertical line is the observed variance of MHC-IIB dissimilarity. The black vertical line is the mean of the simulated distribution, and the black dashed lines are the 2.5 and 97.5 % quantiles of the simulated distribution showing the 95 % confidence intervals. **b-d)** Probability of obtaining a significant result according to the sample size, when considering b) females, and d) males. The dots represent the raw observations from the simulations at each sample size, the line represents the predictions of the model, the shaded area represents the 95 % confidence intervals around the predictions of the model.

**Supporting information 5**: Complementary figures - Probability of divorce (Test 2)

Table S5.1: Statistical significance (column "P") of the relationship between the probability of divorce and the MHC-IIB dissimilarity between the breeding individuals (either females or males, see column "Sex of the individual") and their mates (Test 2). For each model, we reported the "Random effects", and the coefficient estimated by the model as well as their standard error (column "Model estimate") for the associated "Explanatory variable". The maximisation hypothesis and the intermediate optimum hypothesis were tested (column "Hypothesis") for both MHC-IIB functional distance and difference in MHC-IIB supertype sharing (column "MHC dissimilarity index"). See main text for details about the "Probability of significant result" column.

| **Hypothesis** | **MHC dissimila-rity index** | **Sex of the individual** | **Random effects** | **Explanatory variable** | **P** | **Model estimate** | **Odd ratio** | **Probability of significant result** | **Sample size** |
|---|---|---|---|---|---|---|---|---|---|
| - | functional | female | indiv | reproductive outcome | 0.01* | -1.06 ± 0.41 | 0.35 | 0.87 | 1244 |
| intermediate | functional | female | indiv | MHC dissimilarity (quadratic term) | 0.18 | 0.03 ± 0.02 | 1.03 | 0.14 | 1244 |
| maximisation | functional | female | indiv | MHC dissimilarity | 0.58 | -0.08 ± 0.14 | 0.93 | 0.00 | 1244 |
| - | functional | male | indiv, year | reproductive outcome | < 0.01* | -1.50 ± 0.34 | 0.22 | 1.00 | 1232 |
| intermediate | functional | male | indiv, year | MHC dissimilarity (quadratic term) | 0.64 | 0.01 ± 0.03 | 1.01 | 0.02 | 1232 |
| maximisation | functional | male | indiv, year | MHC dissimilarity | 0.68 | 0.03 ± 0.08 | 1.03 | 0.00 | 1232 |
| - | supertype | female | indiv | reproductive outcome | 0.01* | -1.05 ± 0.41 | 0.35 | 0.87 | 1244 |
| intermediate | supertype | female | indiv | MHC dissimilarity (quadratic term) | 0.46 | 0.00 ± 0.00 | 1.00 | 0.05 | 1244 |
| maximisation | supertype | female | indiv | MHC dissimilarity | 0.74 | -0.00 ± 0.01 | 1.00 | 0.00 | 1244 |
| - | supertype | male | indiv, year | reproductive outcome | < 0.01* | -1.50 ± 0.34 | 0.22 | 1.00 | 1232 |
| intermediate | supertype | male | indiv, year | MHC dissimilarity (quadratic term) | 0.81 | -0.00 ± 0.00 | 1.00 | 0.00 | 1232 |
| maximisation | supertype | male | indiv, year | MHC dissimilarity | 0.54 | 0.00 ± 0.01 | 1.00 | 0.01 | 1232 |

*: significance to the level alpha = 0.05. The "-" in any cell of the table shows that the information is not applicable for this column. Note that the sample sizes can be different when considering females and males. For example, the male of a divorcing pair might be observed with a new female and therefore be included in the analyses on divorce probability, while the initial female might not be observed again in the colony.

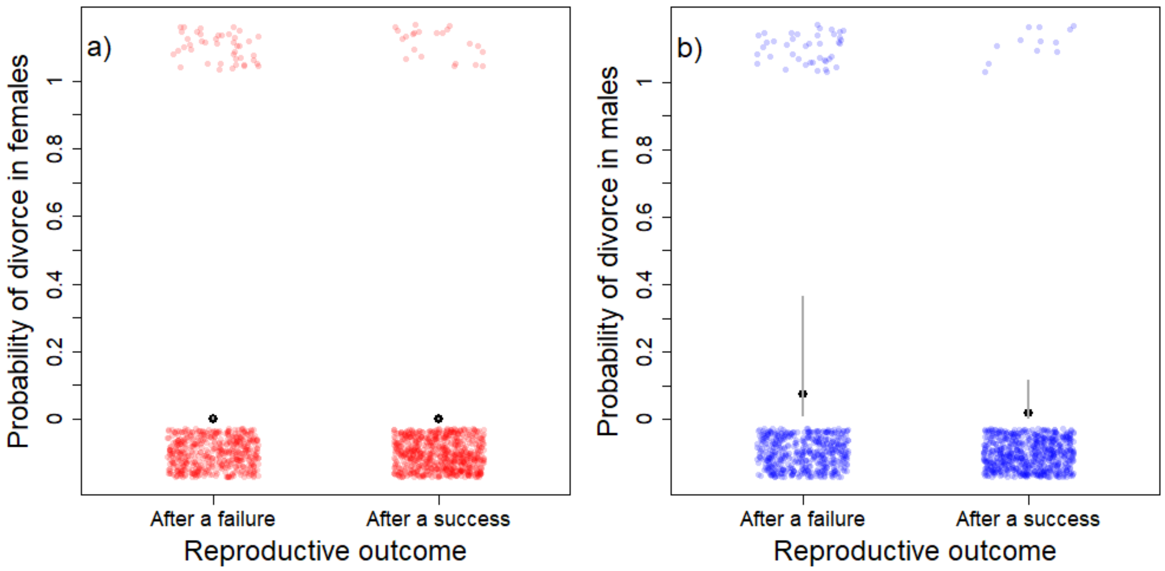


Figure S5.1: Probability of divorce after a reproductive failure and after a reproductive success when considering **a)** females, and **b)** males. The small red and blue dots represent the raw observations, the black large dots represent the predictions of the model. The grey vertical lines represent the 95 % confidence intervals on the fixed effects (note that they are very small when considering females).

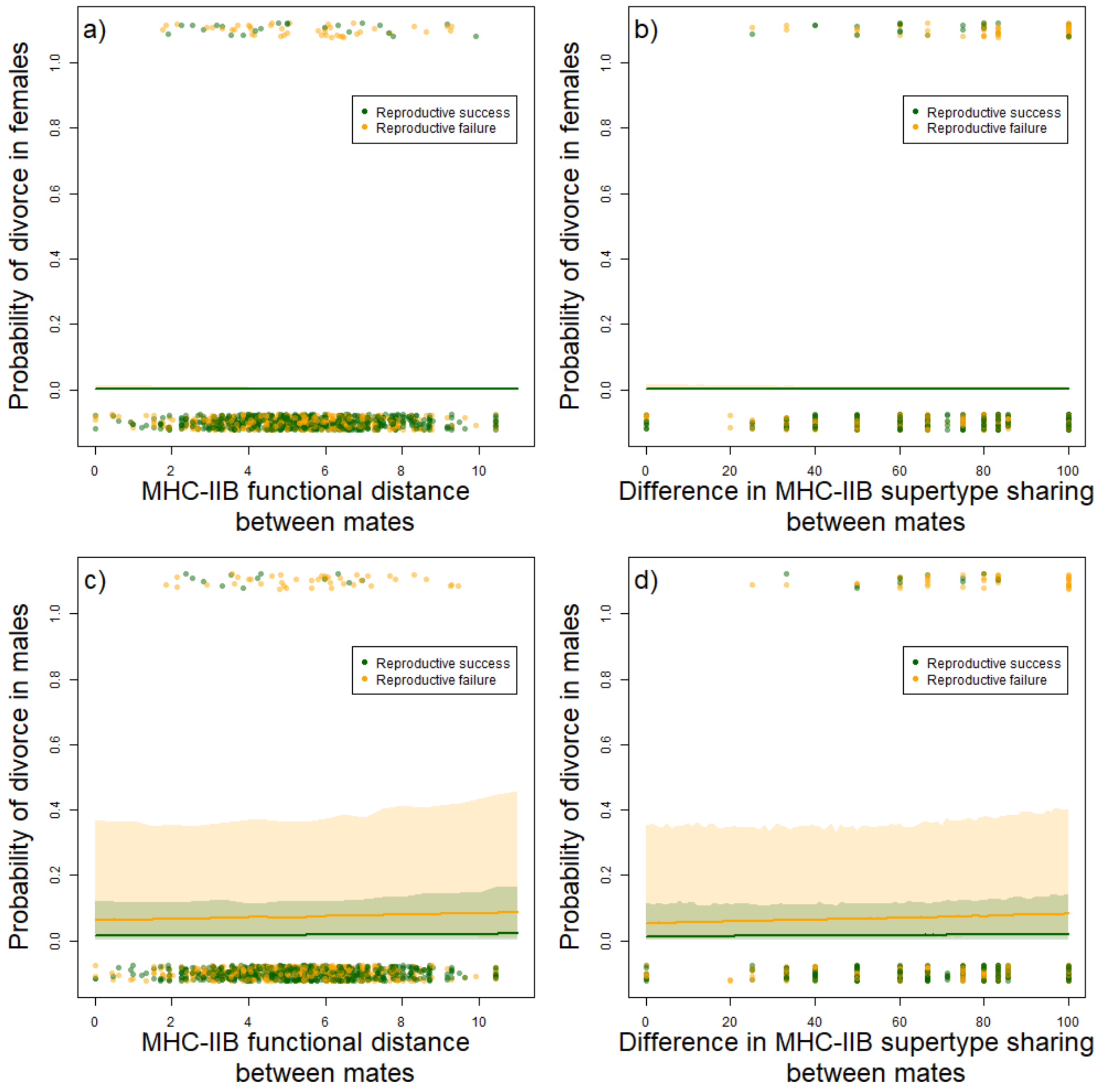


Figure S5.2: Probability of divorce after a reproductive failure (orange) and after a reproductive success (green) when considering a-b) females, and c-d) males, according to **a-c)** the MHC-IIB functional distance with the mates, and **b-d)** the difference in MHC-IIB supertype sharing with the mates. The dots represent the raw observations, the lines represent the predictions of the model, the shaded areas represent the 95 % confidence intervals on the fixed effects.

**Supporting information 6**: Complementary figures - MHC-IIB dissimilarity in original and new pairs after a divorce (Test 3)

Table S6.1: Statistical significance (column “P”) of the variation of MHC-IIB dissimilarity between the breeding individuals (either males or females, see column “Sex of the individual”) and their mates after a divorce (Test 3). The maximisation hypothesis and the intermediate optimum hypothesis were tested (column “Hypothesis”) for both MHC-IIB functional distance and difference in MHC-IIB supertype sharing (column “MHC dissimilarity index”). See main text for details about the “Effect size” column.

| **Hypothesis** | **MHC dissimilarity index** | **Sex of the individual** | **P** | **Effect size** | **Sample size** |
|---|---|---|---|---|---|
| intermediate | functional | female | 0.61 | - | 93 |
| maximisation | functional | female | 0.31 | 0.12 | 93 |
| intermediate | functional | male | 0.66 | - | 65 |
| maximisation | functional | male | 0.67 | 0.07 | 65 |
| intermediate | supertype | female | 0.71 | - | 93 |
| maximisation | supertype | female | 0.36 | 0.11 | 93 |
| intermediate | supertype | male | 0.28 | - | 65 |
| maximisation | supertype | male | 0.59 | 0.09 | 65 |

*: significance to the level alpha = 0.05. The “-” in any cell of the table shows that the information is not applicable for the test.

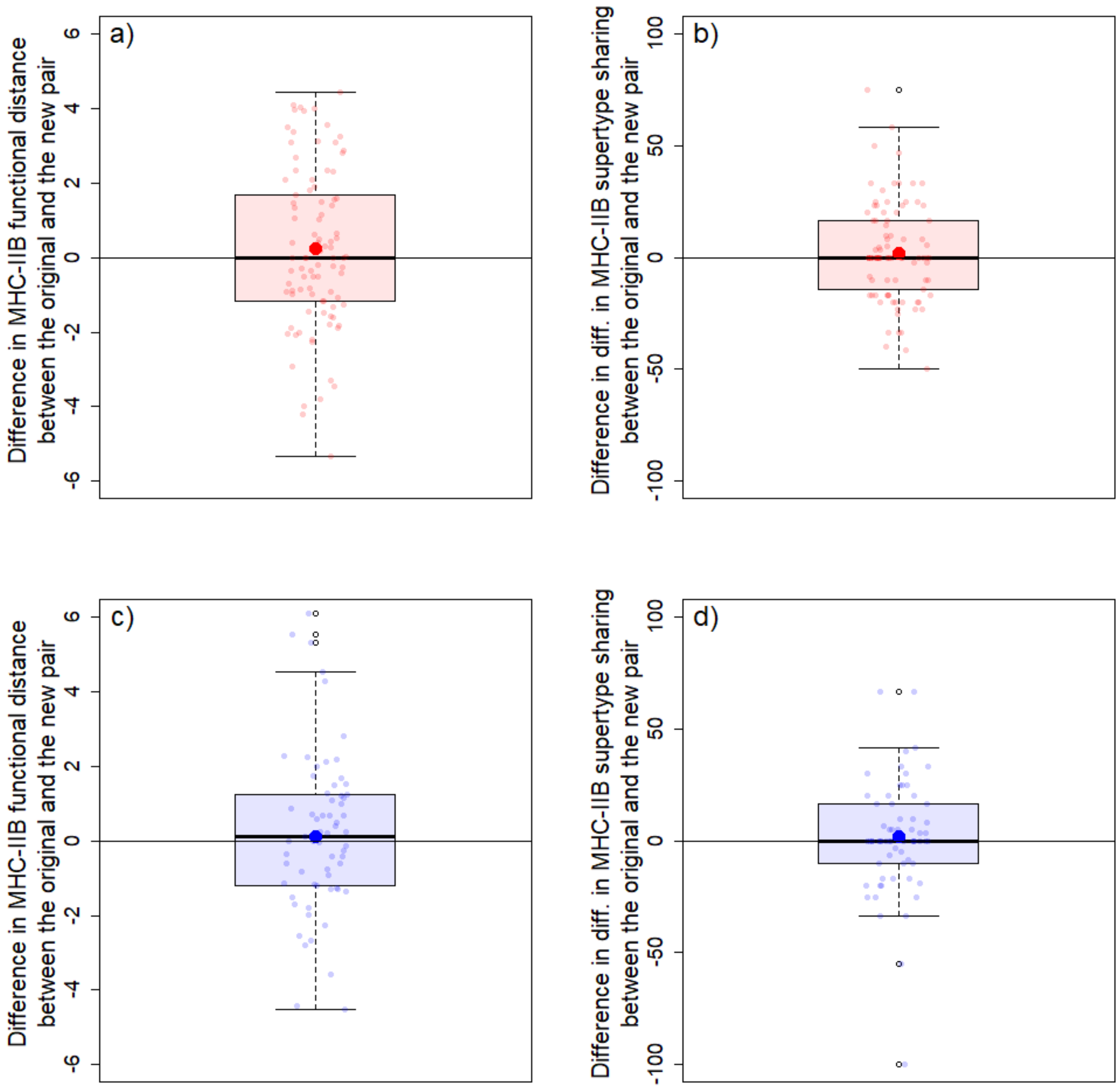


Figure S6.1: Difference in MHC-IIB dissimilarity between the mates in year n+1 (after a divorce) and the mate in year n: **a)** MHC-IIB functional distance when considering females, **b)** difference in MHC-IIB supertype sharing when considering females, **c)** MHC-IIB functional distance when considering males, **d)** difference in MHC-IIB supertype sharing when considering males. The black horizontal line represents a difference in MHC-IIB dissimilarity of 0 between the mates in years n+1 and n.

**Supporting information 7**: Complementary figures - Mate choice in experienced breeders after a divorce (Test 1.2)

Table S7.1: Statistical significance (column "P") of the difference between the observed MHC-IIB dissimilarity (column "Observed value") between the breeding individuals (either females or males, column "Sex of the individual") and their new mates after a divorce, and the random MHC-IIB dissimilarity mean (column "Random estimate", with 95 % confidence intervals) (Test 1.2). The maximisation hypothesis and the intermediate optimum hypothesis were tested (column "Hypothesis") for both MHC-IIB functional distance and difference in MHC-IIB supertype sharing (column "MHC dissimilarity index"). See main text for details about the "Probability of significant result" and "Effect size" columns.

| **Hypothesis** | **MHC dissimila-rity index** | **Sex of the individual** | **P** | **Observed value** | **Random estimate** | **Effect size** | **Probability of significant result** | **Sample size** |
|---|---|---|---|---|---|---|---|---|
| intermediate | functional | female | 0.94 | 3.44 | 3.47 [2.71; 4.32] | -0.03 | 0.00 | 113 |
| maximisation | functional | female | 0.62 | 5.45 | 5.37 [5.06; 5.67] | 0.08 | 0.00 | 113 |
| intermediate | functional | male | 0.79 | 3.39 | 3.27 [2.41; 4.24] | 0.12 | 0.00 | 83 |
| maximisation | functional | male | 0.09 | 5.48 | 5.17 [4.80; 5.53] | 0.31 | 0.21 | 83 |
| intermediate | supertype | female | 0.24 | 359.32 | 438.89 [316.04; 583.88] | -79.57 | 0.00 | 113 |
| maximisation | supertype | female | 0.06 | 77.60 | 74.02 [70.29; 77.67] | 3.58 | 0.23 | 113 |
| intermediate | supertype | male | 0.55 | 380.11 | 428.26 [291.22; 600.64] | -48.15 | 0.00 | 83 |
| maximisation | supertype | male | 0.04* | 76.87 | 72.35 [67.89; 76.52] | 4.52 | 0.78 | 83 |

*: significance to the level alpha = 0.05.

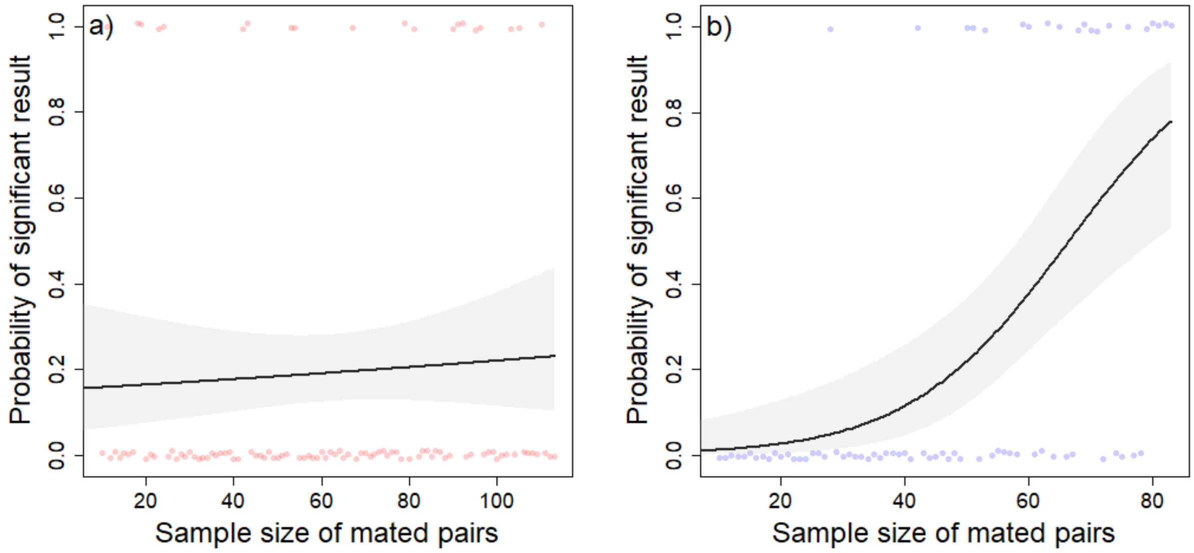


Figure S7.1: Probability of obtaining a significant result according to the sample size when testing the difference in MHC-IIB supertype sharing between divorced breeders and their new mates (cf. Figure 2 in the main text), when considering **a)** females, and **b)** males. The dots represent the raw observations from the simulations at each sample size, the line represents the predictions of the model, the shaded area represents the 95 % confidence intervals around the predictions of the model.

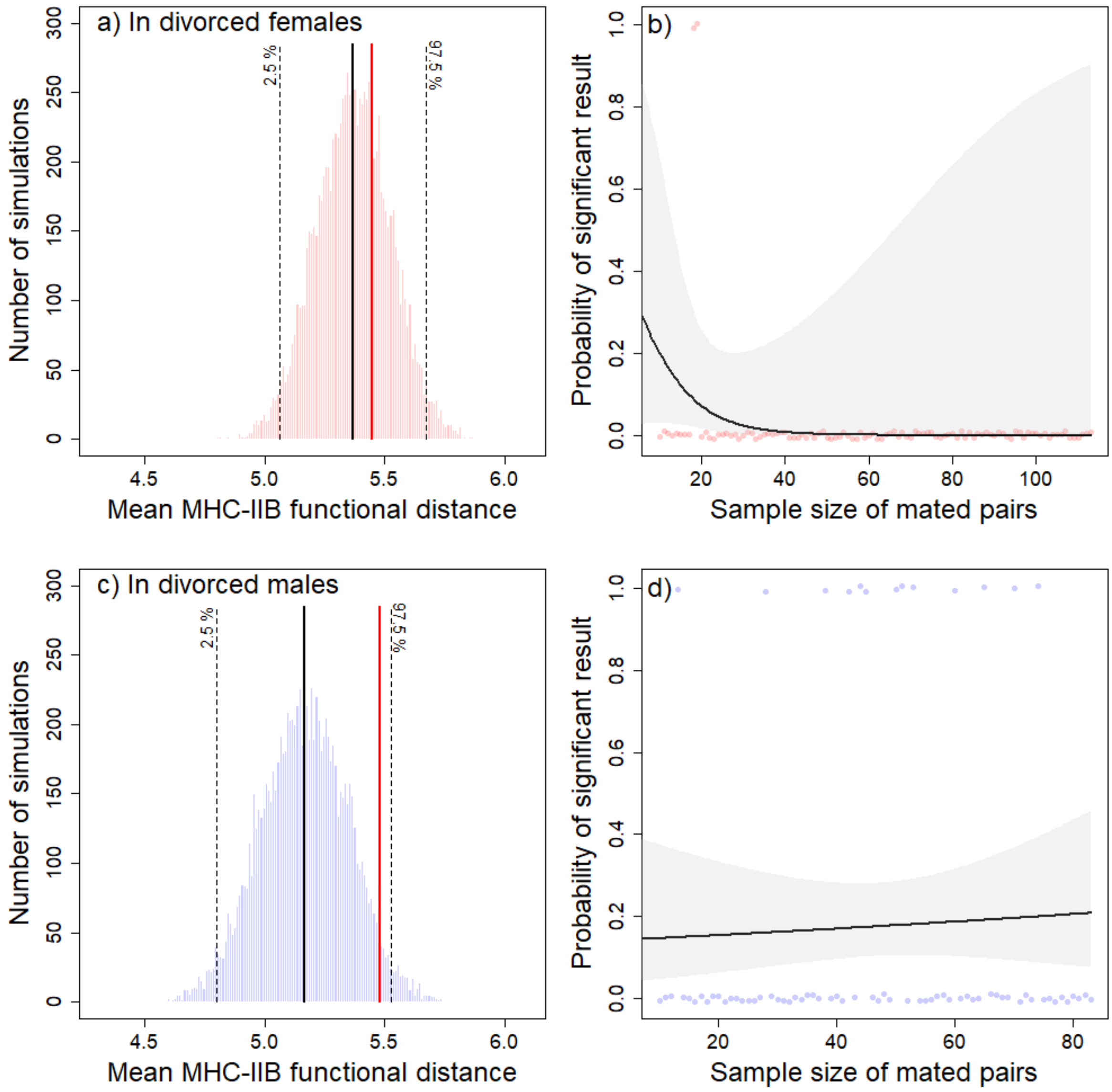


Figure S7.2: **a-c)** Distribution of 10000 means of MHC-IIB dissimilarity (as measured by the MHC-IIB functional distance) between divorced breeders and their new mates, calculated from bootstrap randomisations of potential pairs (see Materials and Methods section for details), when considering a) females, and c) males. The red vertical line is the observed mean of MHC-IIB dissimilarity. The black vertical line is the mean of the simulated distribution, and the black dashed lines are the 2.5 and 97.5 % quantiles of the simulated distribution showing the 95 % confidence intervals. **b-d)** Probability of obtaining a significant result according to the sample size, when considering b) females, and d) males. The dots represent the raw observations from the simulations at each sample size, the line represents the predictions of the model, the shaded area represents the 95 % confidence intervals around the predictions of the model.

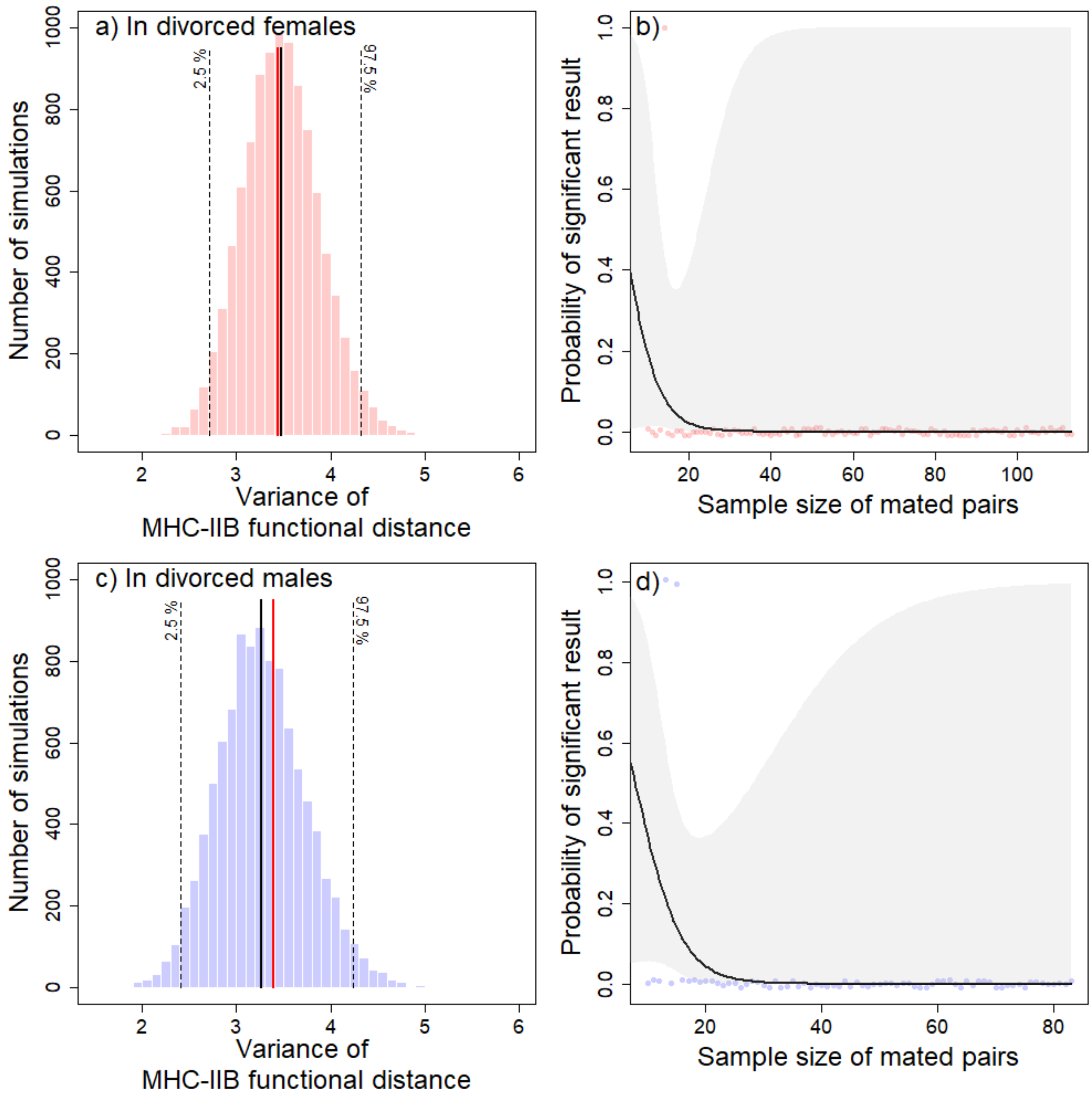


Figure S7.3: **a-c)** Distribution of 10000 variances of MHC-IIB dissimilarity (as measured by the MHC-IIB functional distance) between divorced breeders and their new mates, calculated from bootstrap randomisations of potential pairs (see Materials and Methods section for details), when considering a) females, and c) males. The red vertical line is the observed variance of MHC-IIB dissimilarity. The black vertical line is the mean of the simulated distribution, and the black dashed lines are the 2.5 and 97.5 % quantiles of the simulated distribution showing the 95 % confidence intervals. **b-d)** Probability of obtaining a significant result according to the sample size, when considering b) females, and d) males. The dots represent the raw observations from the simulations at each sample size, the line represents the predictions of the model, the shaded area represents the 95 % confidence intervals around the predictions of the model.

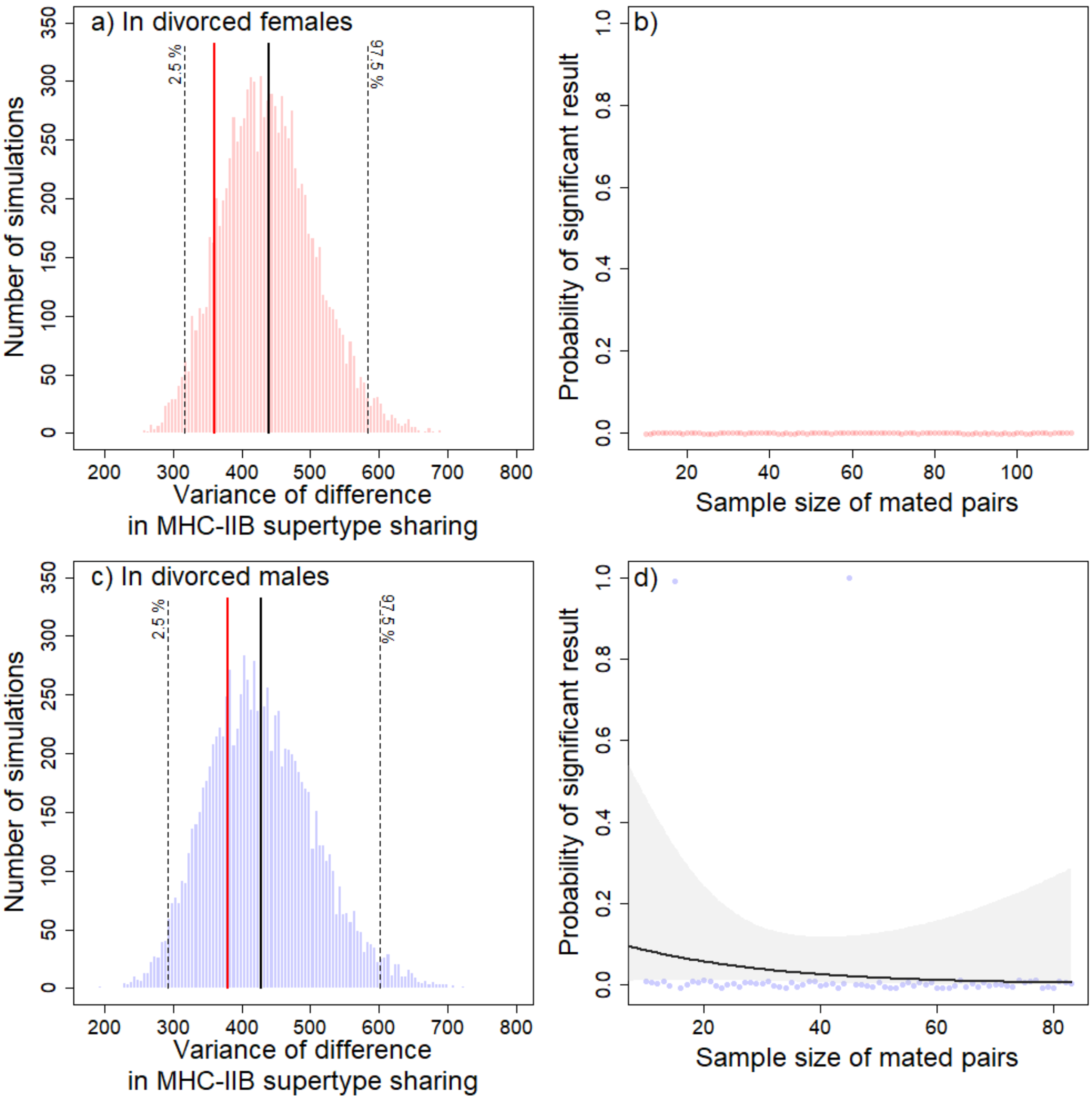


Figure S7.4: **a-c)** Distribution of 10000 variances of MHC-IIB dissimilarity (as measured by the difference in MHC-IIB supertype sharing) between divorced breeders and their new mates, calculated from bootstrap randomisations of potential pairs (see Materials and Methods section for details), when considering a) females, and c) males. The red vertical line is the observed variance of MHC-IIB dissimilarity. The black vertical line is the mean of the simulated distribution, and the black dashed lines are the 2.5 and 97.5 % quantiles of the simulated distribution showing the 95 % confidence intervals. **b-d)** Probability of obtaining a significant result according to the sample size, when considering b) females, and d) males. The dots represent the raw observations from the simulations at each sample size, the line represents the predictions of the model, the shaded area represents the 95 % confidence intervals around the predictions of the model.
